# Social hierarchy assays measure independent features of competitive ability

**DOI:** 10.64898/2026.07.10.737786

**Authors:** Ryan Pitesky, McKinzie Wade, Rachel E. Fanelli, Trevor Rasmuson, Adam C. Nelson, Nicole L. Bedford

## Abstract

In social species, hierarchies emerge across diverse behavioral contexts, including dyadic contests, resource competition, and territorial marking. Whether competitive success generalizes across these contexts remains unclear, in part because individual performance and hierarchy structure are difficult to compare directly across assays. Here, we establish a framework for such comparisons using three behavioral paradigms—the tube test, the warm spot assay, and the void spot assay—in same-sex triads of BALB/c and C57BL/6 mice, as well as *Shank3B* knockouts, which exhibit impaired social recognition. Hierarchies emerged in all three assays, but their structure and behavioral determinants differed by strain and sex. In the tube test, BALB and C57 mice employed different contest-resolution strategies, while *Shank3b*⁻/⁻ females exhibited diminished winner effects and reduced hierarchy stability. In the warm spot assay, females of all strains established stronger hierarchies than males, with high-ranking individuals actively displacing competitors from the warm platform. In the void spot assay, clear hierarchical organization emerged in BALB mice of both sexes and C57 males but was attenuated in *Shank3b*⁻/⁻ mice. Despite repeatable individual performance and stable hierarchies within each assay, performance was weakly correlated between assays, and social rank rarely generalized across behavioral contexts. Together, these findings reveal strain- and sex-dependent strategies for establishing social hierarchies, implicate *Shank3b* in winner effects and hierarchy stability, and demonstrate that competitive ability is multidimensional rather than a single, generalized trait, suggesting that distinct mechanisms may govern competitive success across behavioral domains.

**SIGNIFICANCE STATEMENT:** Social dominance is a fundamental feature of animal societies, but whether the competitive ability that determines rank reflects a stable individual trait or depends on behavioral context remains unclear. We compared social hierarchies across three behavioral paradigms in male and female mice from different genotypes and genetic backgrounds. Using a quantitative framework to compare individual performance and group-level hierarchy structure, we found that competitive success was repeatable within assays but rarely generalized across them. These findings suggest that competitive ability is multidimensional rather than a single, generalized trait. Thus, neural mechanisms identified using one assay may not explain competitive success in other contexts, highlighting the importance of studying social dominance across multiple behavioral domains.

## INTRODUCTION

Social hierarchy—in which the outcome of competitive interactions between individuals becomes predictable—is a pervasive feature of animal groups (1–4). This organizational structure has been documented in insects (5–7), fish (8–10), birds (11–13), rodents (14–16), non-human primates (17–19), and humans (20–22). Across these diverse taxa, position in the hierarchy (i.e., social rank) shapes access to resources such as food, mates, and territory (23) and correlates with physiological state and susceptibility to stress-related disease (24, 25). High-ranking individuals are commonly labeled “dominant” and low-ranking individuals “subordinate”. However, rather than fixed individual traits, these labels may reflect the outcomes of interactions with specific opponents under certain conditions (26). Moreover, individuals may attain high rank through different behavioral strategies (20, 27), and tactics that confer an advantage in one context may be ineffective in another. These observations raise the question of whether social rank generalizes across behavioral domains or, rather, is context-dependent.

If social rank is consistent across domains, competitive ability may reflect a generalized dominance trait, potentially conferred by a shared neural circuit (28). In contrast, if rank is inconsistent across domains, individuals may be dominance specialists, with different traits, and potentially different neural circuits, underlying success in each context (29). By analogy, a top triathlete may perform consistently well in all three events (i.e., generalist) or may excel at one event while performing less well in the others (i.e., specialist). These two routes to success imply different models of how the brain controls competitive performance, which can be distinguished only by tracking the same individuals across multiple domains.

Such comparisons can be challenging because different paradigms may capture distinct aspects of competitive behavior, and hierarchies may be established and expressed differently across assays. Robust cross-domain comparisons must therefore meet three criteria. First, performance must be sufficiently repeatable within each assay to distinguish individual differences from trial-to-trial variation. Second, hierarchy structure must be quantified to determine how competitive ability is distributed within each group. Third, performance measures must be placed on comparable scales to test between-assay consistency and quantify specialization. Together, these measurements provide a framework for separating hierarchy structure from the strategies that produce competitive success and for identifying which aspects of rank, if any, transfer between domains.

Of these criteria, hierarchy structure warrants particular attention because, in groups of three or more individuals, it can reveal opponent-dependent competitive behavior. Specifically, an individual may dominate one group member while submitting to another, a distinction that cannot be established from a single dyadic relationship. Hierarchy type can then be classified by the distribution of competitive ability within the group. For example, a transitive (i.e., linear) hierarchy shows a graded ordering of individuals (A > B > C), whereas a despotic hierarchy is dominated by a single individual (A > B ≈ C). Which type emerges may depend on how well individuals recognize one another. Social recognition—the ability to distinguish individual conspecifics and associate them with the outcomes of previous interactions—may help animals track these relationships and adjust their behavior toward familiar competitors. Such individual-specific information could contribute to establishing and maintaining highly ordered hierarchies (30), yet how social recognition shapes hierarchy structure remains unresolved (3, 31).

Mice (*Mus musculus*) offer a tractable system for addressing whether rank generalizes across domains and what role social recognition plays in hierarchy formation. Mice form social hierarchies both in the wild (32) and in the laboratory (33) and have become an important model organism for investigating the neural mechanisms of social dominance (15). Moreover, mouse hierarchies have been characterized using a variety of behavioral assays (14) and in strains spanning diverse genotypes (34–38) and genetic backgrounds (39, 40).

Here, we studied social hierarchy in both sexes in two strains of laboratory mice commonly used in neurobiological studies (41): BALB/c and C57BL/6 (hereafter BALB and C57). We also included a third strain to test the role of social recognition in hierarchy formation: *Shank3B* knockout mice on a C57 genetic background (hereafter *Shank3b*⁻/⁻) (42). In humans, the *SHANK3* gene is associated with Phelan-McDermid syndrome and autism spectrum disorder and *Shank3b*⁻/⁻ mice show impaired social recognition (42–45), which may alter their capacity to track hierarchical relationships and express rank-appropriate behavior.

We assessed social hierarchy among 68 triads of mice competing in a triathlon of established behavioral paradigms: the tube test, the warm spot assay, and the void spot assay. The tube test is a head-to-head competition in which two mice enter opposite ends of a transparent tube and meet in the middle (46). Because the tube is too narrow for animals to turn around, one must back out. In the warm spot assay, three mice directly compete for access to a warm platform that provides respite from an aversively cold arena (28). In the void spot assay, individual mice respond to a social olfactory cue (i.e., unfamiliar urine) by depositing urinary marks in an open-field arena, signaling social rank (47). By tracking performance across these three paradigms, we assessed how the same individuals organize into social hierarchies in distinct behavioral domains involving dyadic contests, resource competition, and territorial marking.

We set out to compare competitive behavior across multiple domains, characterizing hierarchy structure within each assay and asking whether the consistency of individual rank across domains differs by strain (BALB, C57, *Shank3b*⁻/⁻) and sex. We find that performance is repeatable within a domain, but that dominance (i.e., competitive success) does not readily transfer across domains. Together, these findings indicate that competitive ability is multidimensional, and that the attributes that predict success in one domain are largely independent of those that predict success in another. Thus, rather than reflecting a fixed individual trait governed by a single circuit encoding generalized competitive ability, dominance may arise from multiple domain-specific neural mechanisms.

## RESULTS

### Establishing a comparative framework for social hierarchy across three behavioral domains

Because hierarchies are established and maintained through repeated social interactions (48), mice were co-housed in same-strain, same-sex triads from weaning to ensure familiarity among group members (**Fig. 1A**). To determine whether competitive ability and hierarchy structure generalize across domains (**Fig. 1B**), we tested mice in three behavioral paradigms: the tube test, the warm spot assay, and the void spot assay. For the tube test, mice were trained over the course of a week to traverse the tube freely in both directions (**Fig. 1C**). Triads then completed a 30-trial round-robin tournament in which every mouse encountered each of its two opponents 10 times. Trial duration and resolution mode (winner advance or loser retreat) were recorded for all trials. For the warm spot assay, all three mice were simultaneously placed on a cold plate (15°C) with access to a small warm platform (25°C) (**Fig. 1D**). Each triad was tested twice in 20-min trials, and individual warm spot bouts were scored blind to experimental condition (i.e., strain and sex). For the void spot assay, individual mice were placed in an open-field arena lined with filter paper and allowed to freely explore for 20 min (**Fig. 1E**). Two baseline trials without stimulus were followed by two trials in which 60 µL of unfamiliar same-sex urine was applied to the center of the filter paper. The resulting urinary void spots were imaged and quantified to extract spot number, volume, and spatial distribution across the arena (49). All assays were conducted during the dark phase and video-recorded under infrared illumination. See Extended Methods for full experimental details.

**Figure 1.**
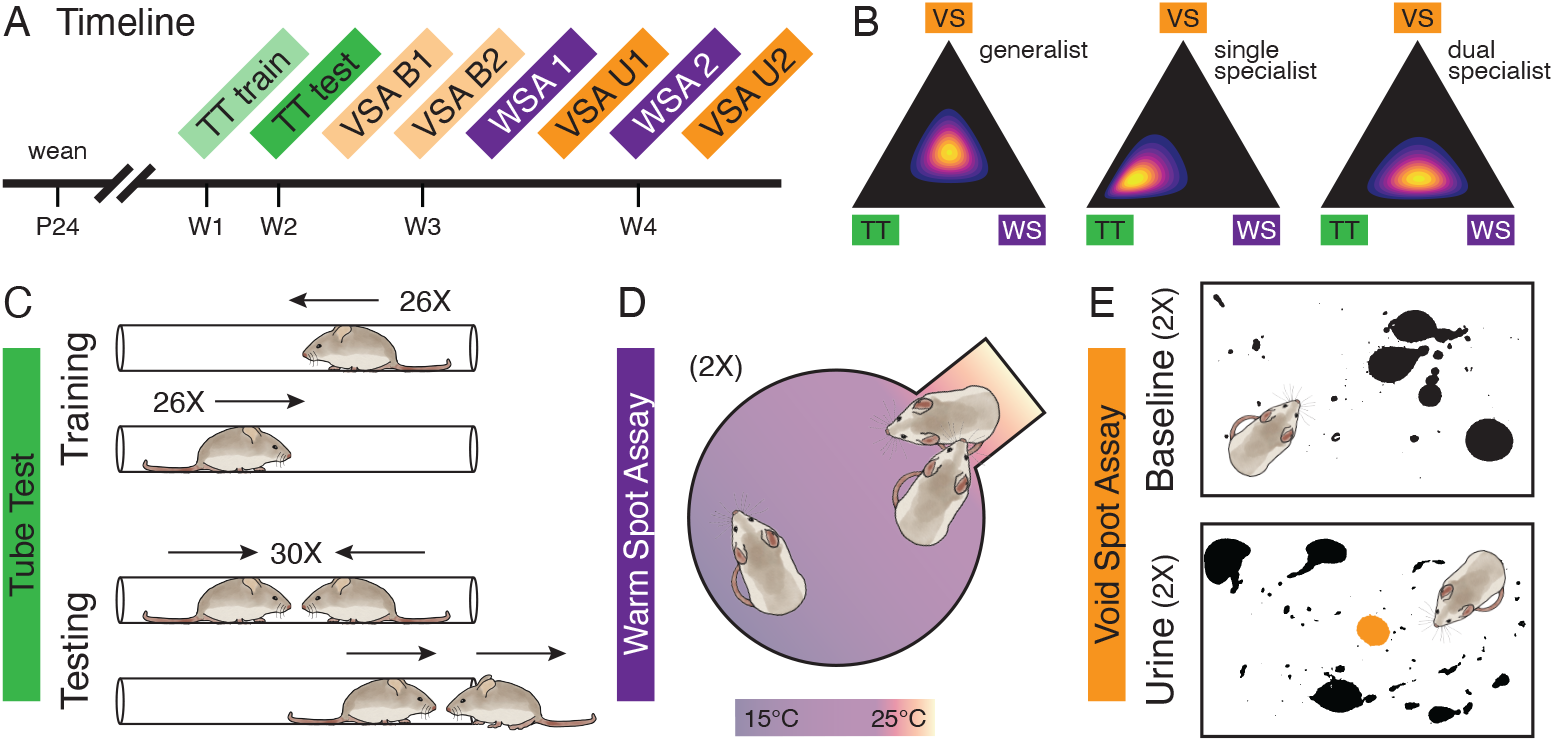
Experimental timeline, predictions, and behavioral assay design. **(A)** Age-matched mice from different litters were weaned at postnatal day 24 (P24) into same-sex groups of three. Co-housed triads then underwent 4 weeks of behavioral testing. **(B)** Simulated ternary plots illustrating generalist and specialist patterns of behavioral performance. The center of the plot represents balanced performance across assays, consistent with a generalist strategy. Displacement toward a vertex represents greater specialization in a single assay (single specialist), whereas displacement toward an edge represents greater specialization in two assays (dual specialist). **(C)** Tube test (TT) training and testing schematic. Mice were trained to traverse the tube freely 26 times in each direction. During testing, each mouse completed 10 trials against each opponent for a total of 30 trials per triad. **(D)** Warm spot assay (WSA) schematic. Mice were introduced to a 15°C cold plate with access to a 25°C warm spot. Each triad received two 20-min trials. **(E)** Void spot assay (VSA) schematic. Individual mice were introduced to an open-field arena lined with filter paper for 20 min, either without urine stimulus (baseline) or with 60 µL of same-sex urine pipetted onto the center of the paper. Each mouse received two baseline (B) and two urine stimulus (U) trials. All behavioral assays were conducted during the dark phase.

### Tube test hierarchies exhibit strain- and sex-dependent organization and are disrupted by *Shank3b* mutation

#### Trial Duration and Resolution Mode

To determine whether tube test behavior differed by mouse strain or sex, we first compared trial duration and resolution mode across groups. The duration of tube test contests differed considerably among strains (**Fig. 2A**). Specifically, Tukey-adjusted post hoc contrasts from a linear mixed-effects model (LMM) confirmed that BALB mice took significantly longer to resolve contests (median [IQR] = 51 s [27.8–84.2 s]) than either C57 (t = 10.54, P < 0.001) or *Shank3b*⁻/⁻ mice (t = 5.62, P < 0.001). Contest duration also differed significantly between C57 and *Shank3b*⁻/⁻ (t = 3.09, P = 0.007), with C57 mice resolving contests twice as fast (12 s [7–20 s]) as *Shank3b*⁻/⁻ mice (24 s [15–40.8 s]). In addition, BALB opponent pairs more frequently resolved contests by the winner advancing rather than the loser retreating, compared with both C57 (t = 2.68, P = 0.022) and *Shank3b*⁻/⁻ pairs (t = 3.39, P = 0.003) (**Fig. 2B**). Together, these findings indicate that (1) BALB mice take longer to complete tube tests than C57 mice and resolve them differently, and (2) *Shank3b⁻/⁻* mice have longer decision times than their C57 counterparts.

**Figure 2.**
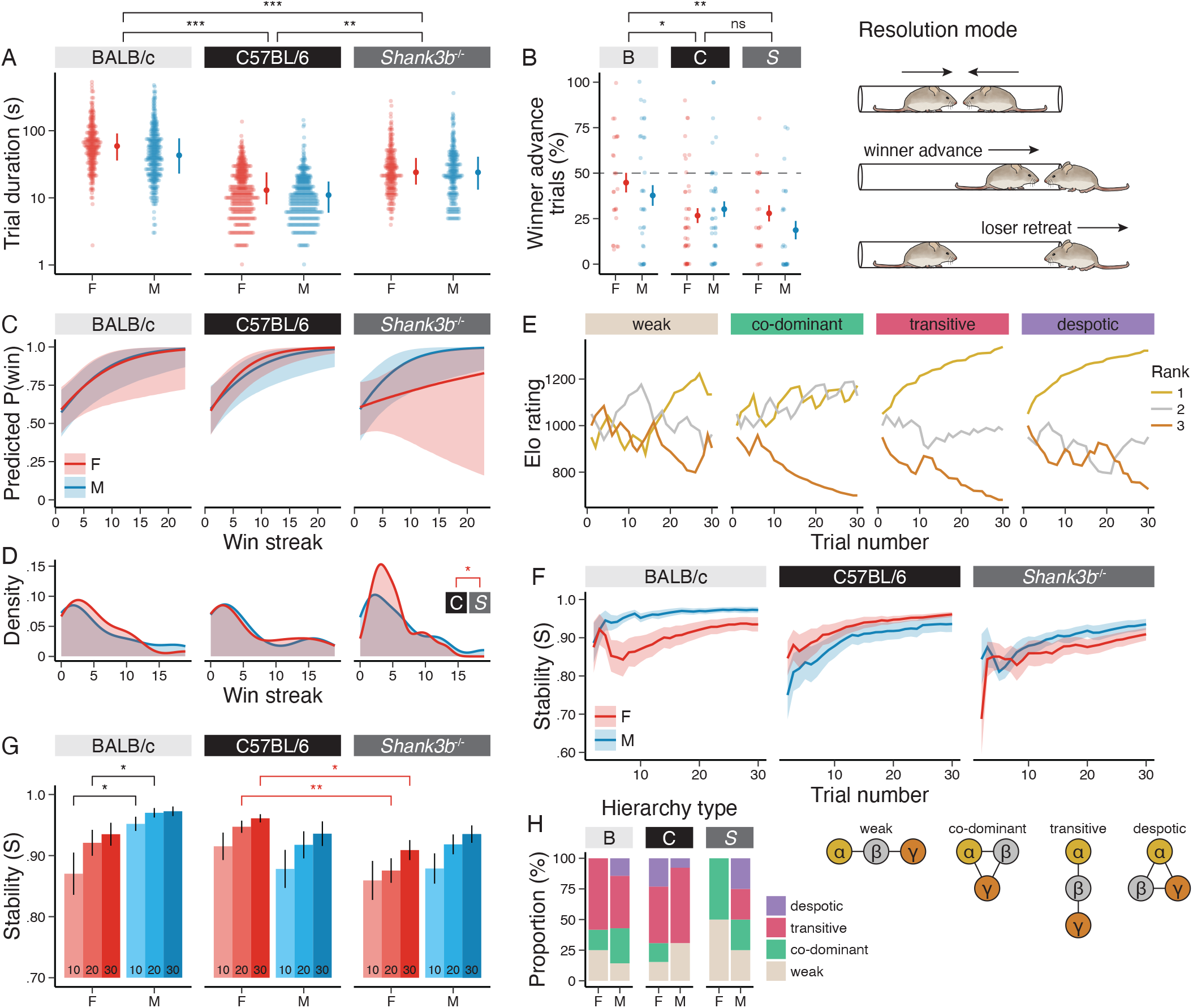
Variation in tube test behavior among mouse strains. **(A)** Trial duration by strain and sex. Points indicate median duration and bars indicate the interquartile range (IQR). **(B)** Proportion of trials resolved by winner advancing vs. loser retreating. Points indicate mean proportion per opponent pair and bars indicate ± SE. **(C)** Posterior predicted probability of winning as a function of win streak length. Shaded areas denote 95% credible intervals. **(D)** Maximum win streak length distributions. **(E)** Representative plots showing change in Elo rating over time. **(F)** Change in hierarchy stability (S index) over time. Lines indicate means and shaded areas denote ± SE. **(G)** Hierarchy stability at trials 10, 20, and 30. Bar heights indicate means and error bars indicate ± SE. **(H)** Proportion of hierarchy types (determined by David’s score) by strain and sex. Abbreviations: F = female; M = male; B = BALB/c; C = C57BL/6; S = *Shank3b*^-/-^; α = rank 1; β = rank 2; γ = rank 3.

#### Winner Effects

Winner effects, whereby previous contest experience affects the outcome of future interactions, can play an important role in shaping hierarchies (50). To determine how prior wins affect future wins across strains and sexes, we fit a Bayesian mixed-effects logistic regression model predicting the probability of winning as a function of win streak length, with random intercepts for focal mouse and partner identity. Overall, win streak length positively predicted contest outcome, with each additional prior win increasing the odds of winning the next contest by 19% (odds ratio = 1.19, 95% credible interval [1.03, 1.39]). Estimated marginal slopes indicated consistent winner effects across all strains and both sexes, except in *Shank3b*⁻/⁻ females, where the slope was weak and not credibly different from zero (**Fig. 2C**). Additionally, the distribution of maximum win streak lengths differed significantly between C57 and *Shank3b*⁻/⁻ females (Anderson–Darling test: AD = 2.47, P = 0.049), reflecting shorter win streaks in mutant (mean ± SE = 4.96 ± 0.65) than in wild-type females (6.15 ± 1.08) (**Fig. 2D**). These results indicate that winner effects are diminished in *Shank3b*⁻/⁻ females.

#### Hierarchy Stability

To compare hierarchy formation dynamics across strains and sexes, we used Elo ratings (51), which are updated after each contest based on expected vs. observed outcomes, with unexpected outcomes resulting in larger rating shifts (**Fig. 2E**). We then examined hierarchy stability (i.e., consistency in the ordinal rank of Elo ratings over time) using the S index (52). Across strains and sexes, hierarchies became more stable over time (LMM: t = 6.82, P < 0.001) (**Fig. 2F**). Within BALB, male triads were more stable than female triads at trial 10 (t = 2.21, P = 0.031) and trial 20 (t = 2.05, P = 0.045), but this sex difference was attenuated by trial 30 (t = 1.79, P = 0.079) (**Fig. 2G**). Additionally, although C57 and *Shank3b*⁻/⁻ female triads were similarly stable at trial 10 (t = 1.87, P = 0.158), mutant females were significantly less stable than their wild-type counterparts at both trial 20 (t = 3.17, P = 0.007) and trial 30 (t = 2.62, P = 0.030). In contrast, C57 and *Shank3b*⁻/⁻ male triads showed similar stability trajectories over time. These findings suggest that *Shank3b⁻/⁻* females have a reduced capacity to establish stable hierarchies in head-to-head competition.

#### Hierarchy Structure

Cyclic dominance, in which individuals dominate one group member but submit to another in a rock–paper–scissors pattern, was rare, occurring in only 2 of 68 triads. To characterize hierarchy structure, we classified each triad based on the magnitude and distribution of differences in competitive ability among group members. First, as a measure of competitive ability in the tube test, we calculated David’s score for each mouse, which is derived from the proportion of wins and losses against each opponent, weighted by the relative dominance of those opponents (53). Next, we categorized each hierarchy based on the spread of normalized David’s scores within each triad (see Extended Methods). Triad hierarchies were classified as “weak” (i.e., egalitarian) if the difference in normalized David’s score between the rank 1 and rank 3 individual fell within the bottom quartile of the distribution across all triads, regardless of strain or sex; all remaining triads were classified as hierarchical. Hierarchical triads were further subcategorized as “co-dominant” when rank 2 was closer to rank 1, “transitive” when rank 2 was centrally positioned between ranks 1 and 3, and “despotic” when rank 2 was closer to rank 3. The distribution of hierarchy types differed significantly among strains in females (Fisher’s exact test: P = 0.024) but not males (P = 0.259) (**Fig. 2H**). Pairwise comparisons in females revealed that the distribution of hierarchy types in *Shank3b⁻/⁻* triads differed from that of both C57 (P = 0.014) and BALB triads (P = 0.028), whereas C57 and BALB triads did not differ from one another (P = 0.456). This pattern was driven by (1) a higher prevalence of weak hierarchies among *Shank3b⁻/⁻* females (50%) than among wild-type (i.e., BALB and C57) females (20% on average), and (2) the absence of transitive and despotic hierarchies among mutant females. These results suggest reduced hierarchical organization in females with impaired social recognition.

#### Tube Test Summary

Overall, BALB mice of both sexes took nearly three times as long to resolve tube test contests and were more likely than mice on a C57 background to resolve contests by the winner pushing out the loser. Nevertheless, BALB mice (particularly males) achieved a high degree of hierarchy stability, indicating that slow contest resolution does not preclude the formation of stable dominance hierarchies. Among mice on a C57 background, wild-type mice resolved contests twice as fast as *Shank3b*⁻/⁻ mice. In addition, winner effects were diminished in *Shank3b*⁻/⁻ females, which accumulated shorter win streaks than C57 females. *Shank3b*⁻/⁻ female hierarchies were also less stable than those of C57 females, with more frequent rank reversals throughout the round-robin tournament, consistent with the higher proportion of weak hierarchies observed in this group. In summary, these findings indicate that, while all mice established stable hierarchies in the tube test, the temporal dynamics, strength, and structure of those hierarchies differed by strain and sex.

### Warm spot assay reveals strain-dependent displacement strategies and despotic hierarchies in female triads

#### Warm Spot Duration

First, to confirm that the warm spot is a desirable resource worth competing for, we measured total platform occupancy. Summing the total durations of all three mice, we found a cumulative warm spot occupancy of 19.7 ± 0.1 min (mean ± SE) per 20-min trial. However, across all strains and both sexes, warm spot occupancy differed among individuals within each triad, resulting in a consistent rank order based on cumulative warm spot duration (**Fig. 3B**). Overall, females spent more time on the warm spot than males (t = 4.84, P < 0.001), and *Shank3b*⁻/⁻ mice spent more time on the warm spot than either C57 (t = 5.23, P < 0.001) or BALB mice (t = 3.67, P < 0.001). This effect was largely driven by females, with *Shank3b*⁻/⁻ females spending more time on the warm spot than either C57 (t = 3.93, P < 0.001) or BALB females (t = 3.39, P = 0.003), while *Shank3b*⁻/⁻ males showed greater cumulative warm spot duration than C57 (t = 3.68, P < 0.001) but not BALB males (t = 2.03, P = 0.109) (**Fig. 3C**). Together, these findings suggest that the warm spot assay captures individual differences in competitive access to a desirable, limited resource.

**Figure 3.**
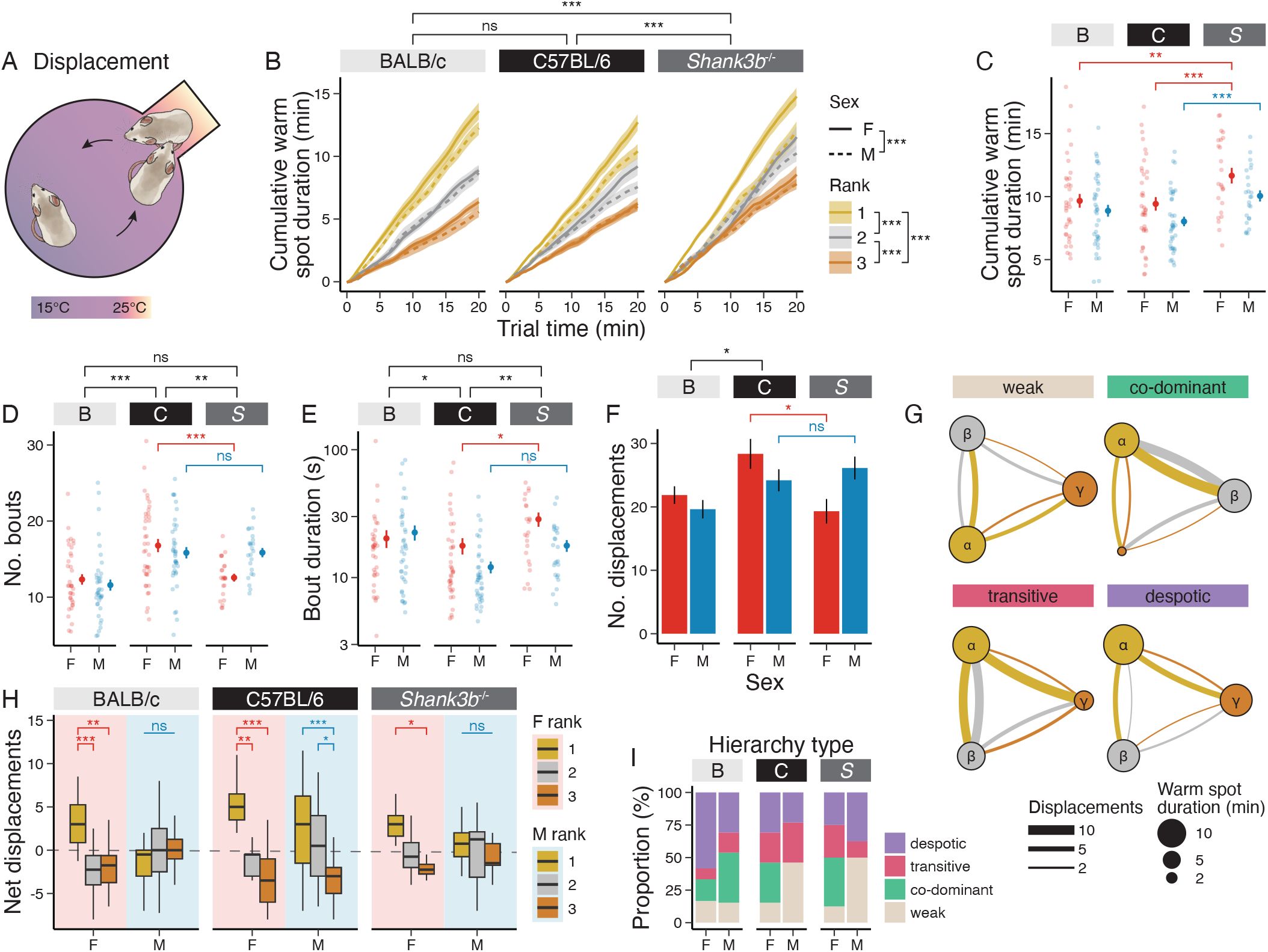
Variation in warm spot behavior among mouse strains. **(A)** Schematic of the warm spot assay illustrating displacement behavior. **(B)** Cumulative warm spot duration by strain, sex, and rank. Ranks are based on cumulative warm spot duration (rank 1 = most time, rank 3 = least time). Solid lines represent females and dashed lines represent males. Lines indicate means and shaded areas denote ± SE. **(C)** Mean cumulative warm spot duration per mouse. **(D)** Mean number of warm spot bouts per mouse. **(E)** Median bout duration per mouse. For (C), (D), and (E), each point represents the mean/median value per mouse across two trials and error bars denote ± SE. **(F)** Total number of displacements per trial per triad. Bar heights indicate means and error bars indicate ± SE. **(G)** Representative network graphs with vertex size scaled by cumulative warm spot duration and edge weight scaled by number of displacements. **(H)** Net displacements by strain, sex, and rank. **(I)** Proportion of hierarchy types (determined by cumulative warm spot duration) by strain and sex.

#### Warm Spot Dynamics

Next, to determine how warm spot occupancy dynamics vary across strains and sexes, we quantified the number of warm spot bouts and the median bout duration for each mouse. Overall, C57 mice had more bouts than either BALB (t = 5.37, P < 0.001) or *Shank3b*⁻/⁻ mice (t = 3.35, P = 0.003) (**Fig. 3D**). Notably, Tukey-adjusted post hoc contrasts revealed that the difference between C57 and *Shank3b*⁻/⁻ mice was present in females (t = 4.28, P < 0.001) but not males (t = 0.57, P = 0.835). C57 mice also had shorter bout durations than both BALB (t = 2.37, P = 0.049) and *Shank3b*⁻/⁻ mice (t = 3.10, P = 0.006). Again, the difference between C57 and *Shank3b*⁻/⁻ mice was evident in females (t = 2.87, P = 0.013) but not males (t = 1.64, P = 0.233) (**Fig. 3E**). Thus, C57 mice occupied the warm spot in more frequent but shorter bouts than BALB and *Shank3b⁻/⁻* mice, with the effect of *Shank3b* mutation specific to females.

#### Displacements

To assess agonistic interactions in the warm spot assay, we quantified displacement behavior (i.e., how often a mouse forced another off the platform). Overall, BALB triads had fewer total displacements per trial than C57 triads (t = 2.90, P = 0.015) (**Fig. 3F**). In addition, *Shank3b*⁻/⁻ female triads (t = 3.24, P = 0.005), but not male triads (t = 0.10, P = 0.995), showed significantly fewer displacements than their C57 counterparts. These findings suggest that *Shank3b*⁻/⁻ females show resource competition behaviors more similar to BALB mice (fewer, longer bouts and fewer displacements), whereas *Shank3b*⁻/⁻ males more closely resemble C57 mice (more frequent, shorter bouts and more displacements). These results raise the possibility that impaired social recognition influences the dynamics of resource competition in a sex-dependent manner.

Next, we examined the relationship between rank (determined by mean cumulative warm spot duration) and net displacements (i.e., the number of times a mouse displaced another minus the number of times it was displaced) (**Fig. 3G**). Overall, rank 1 females had higher net displacement values than both rank 2 (BALB: t = 4.15, P < 0.001; C57: t = 3.64, P = 0.001; *Shank3b*⁻/⁻: t = 1.70, P = 0.210) and rank 3 females (BALB: t = 3.45, P = 0.002; C57: t = 5.37, P < 0.001; *Shank3b*⁻/⁻: t = 2.78, P = 0.017) (**Fig. 3H**). C57 males showed a similar pattern: rank 1 (t = 3.76, P < 0.001) and rank 2 males (t = 2.58, P = 0.029) had higher net displacement values than rank 3 males. In contrast, there was no significant effect of rank on displacement behavior in either BALB or *Shank3b*⁻/⁻ males. Together, these findings suggest that higher-ranking females of all strains tend to monopolize the warm spot by displacing other mice—a pattern also present in C57 males but absent in BALB and *Shank3b*⁻/⁻ males.

#### Hierarchy Structure

We assigned hierarchy types based on the spread of cumulative warm spot duration values (averaged across two trials) within each triad. While there was no sex difference in BALB mice (Fisher’s exact test: P = 0.529), among mice on a C57 background, the distribution of hierarchy types differed significantly between sexes (P = 0.009) (**Fig. 3I**).

Combining C57 and *Shank3b*⁻/⁻ triads, we found that weak hierarchies were more prevalent in males (48%) than females (14%), suggesting that competition over access to warmth produces more pronounced rank differentiation in females. These results indicate that sex differences in warm spot hierarchy structure depend on genetic background, emerging in C57-background mice but not in BALB.

#### Warm Spot Summary

Altogether, these findings reveal that similar levels of resource occupancy can arise through distinct behavioral dynamics. Although BALB and C57 mice spent comparable amounts of time on the warm spot, BALB mice had fewer, longer bouts and fewer displacements, whereas C57 mice had more frequent, shorter bouts and higher displacement rates. Notably, *Shank3b*⁻/⁻ females resembled BALB rather than C57 females in bout structure and displacement frequency, whereas *Shank3b*⁻/⁻ males exhibited patterns more similar to their C57 counterparts. Finally, hierarchy structure varied by sex and strain, with strong hierarchies predominating in females and BALB males, but weak hierarchies predominating in C57 and *Shank3b*⁻/⁻ males.

### Void spot assay reveals strong hierarchical organization in BALB mice and C57 males but attenuated structure in *Shank3b⁻/⁻* mice

#### Void Frequency and Volume

To assess how performance in urinary marking behavior varies across strains and sexes in response to a social olfactory cue, we quantified void frequency and volume in the presence and absence of an unfamiliar urine stimulus. We assessed voiding behavior using a hurdle modeling approach, first analyzing whether animals voided (binary outcome), then comparing total void volume among trials in which voiding occurred. Both baseline (B) and urine stimulus (U) trials were included in the analysis. Tukey-adjusted post hoc contrasts from a generalized linear mixed-effects model (GLMM) indicated that BALB mice were more likely to void than either C57 (z = 8.22, P < 0.001) or *Shank3b*⁻/⁻ mice (z = 7.09, P < 0.001) (**Fig. 4A**). While all three strains were more likely to void in response to a urine stimulus (z = 9.29, P < 0.001), BALB mice showed only a slight increase, whereas mice on a C57 background approximately doubled their response rates. In trials where voiding occurred, BALB mice had higher total void volumes than either C57 (t = 10.01, P < 0.001) or *Shank3b*⁻/⁻ mice (t = 9.18, P < 0.001), while C57 and *Shank3b*⁻/⁻ mice were statistically indistinguishable (t = 1.00, P = 0.576). Tukey-adjusted post hoc contrasts confirmed that males voided more than females in both wild-type strains (BALB: t = 3.65, P < 0.001; C57: t = 6.49, P < 0.001), whereas no significant sex differences were observed in *Shank3b*⁻/⁻ mice (t = 1.88, P = 0.061) (**Fig. 4B**). Together, these findings demonstrate that although *Shank3b*⁻/⁻ mice retain a robust urinary response to unfamiliar social cues, they lack the pronounced sex difference in void volume observed in both wild-type strains.

**Figure 4.**
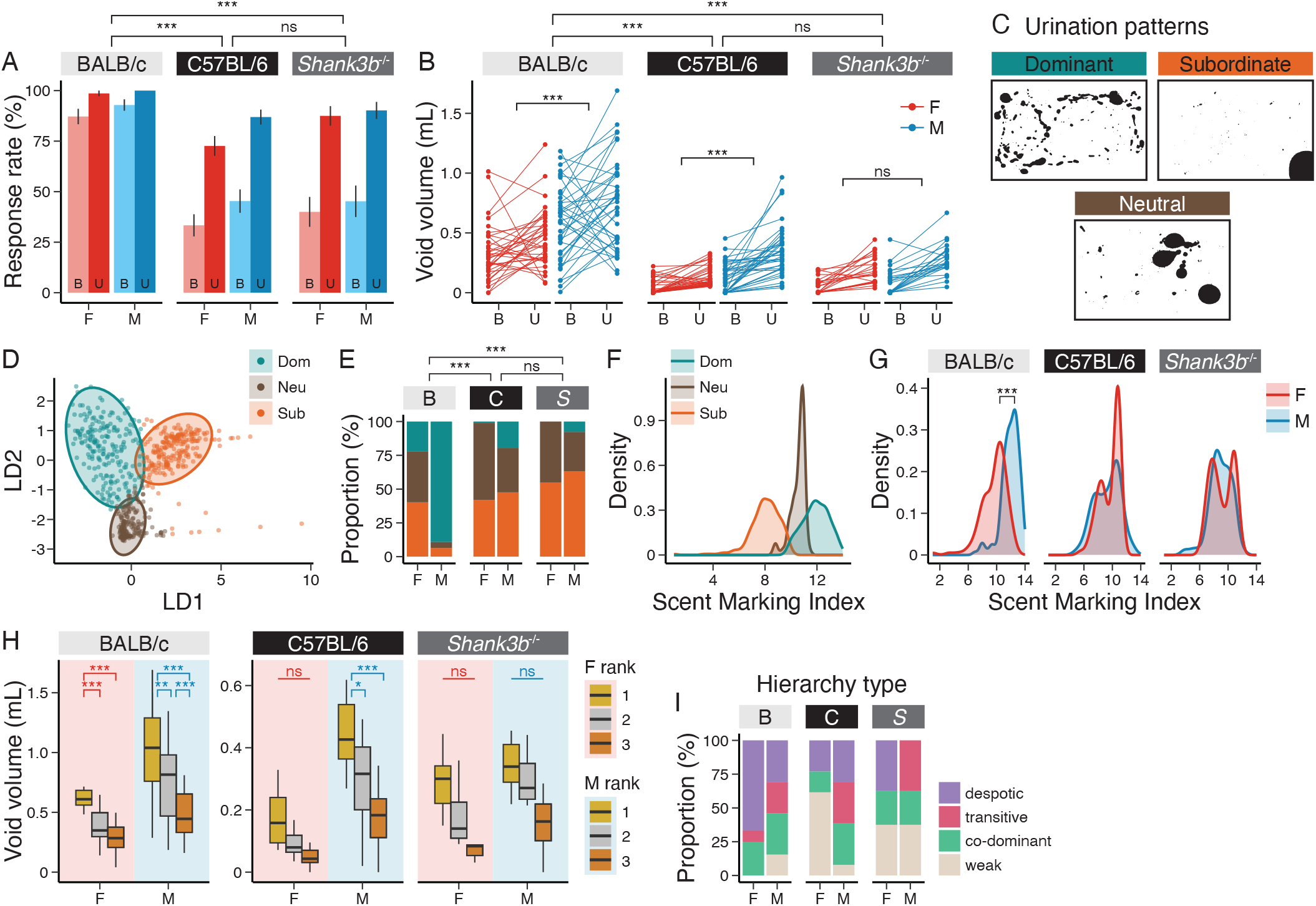
Variation in urination behavior among mouse strains. **(A)** Probability of voiding in baseline (B) and urine stimulus (U) trials by strain and sex. Bar heights indicate means and error bars indicate ± SE. **(B)** Total void volume by strain and sex. Each point represents the mean of two baseline (B) or two urine stimulus (U) trials per mouse. **(C)** Representative urination pattern classifications. **(D)** Projection of urination patterns onto the first two linear discriminant axes (LD1 and LD2). Each point represents a trial in which voiding occurred, and ellipses indicate the spread of each predicted class. **(E)** Distribution of urination pattern classes by strain and sex. **(F)** Scent marking index (SMI) distributions by urination pattern. **(G)** SMI distributions by strain and sex. **(H)** Total void volume by strain, sex, and rank. **(I)** Proportion of hierarchy types (determined by total void volume) by strain and sex.

#### Urination Pattern Classification

We used linear discriminant analysis (LDA) to classify urination patterns as dominant, subordinate, or neutral (**Fig. 4C**; see Extended Methods for definitions). Clear distinctions were observed among the three urination patterns, with cross-validated sensitivities of 91%, 87%, and 89% for dominant, subordinate, and neutral classes, respectively. The first linear discriminant axis (LD1) separated dominant from subordinate patterns, whereas LD2 distinguished neutral from the other classes (**Fig. 4D**). We found significant differences in the distribution of urination pattern classes across strains and sexes (**Fig. 4E**). Fisher’s exact tests revealed that BALB differed significantly from both C57 (P < 0.001) and *Shank3b*⁻/⁻ (P < 0.001). For example, dominant urination patterns were observed in 22.3% of BALB female trials, compared with only 1.2% of C57 female trials; no *Shank3b*⁻/⁻ females exhibited dominant urination behavior. Among males, dominant urination patterns were observed in 89.1% of BALB trials, compared with only 19.6% and 7.7% of C57 and *Shank3b*⁻/⁻ trials, respectively.

Next, we derived a scent marking index (SMI) by inverting and shifting LD1 so that the minimum SMI = 1, with higher values indicating more dominant-like urination patterns (**Fig. 4F**). BALB mice had higher SMI values than both C57 (t = 8.23, P < 0.001) and *Shank3b*⁻/⁻ mice (t = 8.20, P < 0.001) (**Fig. 4G**). BALB males also had higher SMI values than BALB females (t = 9.75, P < 0.001), whereas no sex differences were observed in either C57 (t = 0.98, P = 0.331) or *Shank3b*⁻/⁻ mice (t = 0.70, P = 0.485). Together, these results reveal strain and sex effects on marking behavior. Dominant-like marking was more prevalent in BALB mice than in C57-background mice, and dominant urination patterns were more common in males than females across strains, although sex differences in SMI were significant only in BALB mice. *Shank3b*⁻/⁻ mice showed the lowest incidence of dominant urination patterns, which were absent in females.

#### Void Volume by Rank

Although SMI captures broad differences in urination patterns across strains and sexes, multiple individuals within a triad may exhibit similar patterns (e.g., dominant-like), making it difficult to distinguish their relative ranks. To assess rank differentiation within triads, we therefore ranked mice by total void volume (averaged across two urine stimulus trials) and compared these values among ranks (**Fig. 4H**). BALB triads of both sexes exhibited clear hierarchical organization, with significant separation between ranks (rank 1 vs. rank 3: t = 8.99, P < 0.001), as did C57 males (t = 4.44, P < 0.001). In contrast, hierarchical organization was attenuated in C57 females and in *Shank3b⁻/⁻* triads of both sexes. The presence of rank-differentiated urination behavior in BALB females is consistent with recent evidence that BALB females engage in territorial scent marking (54).

#### Hierarchy Structure

We then classified hierarchy type based on the spread of total void volumes (averaged across two urine stimulus trials) within each triad. The distribution of hierarchy types differed significantly among strains in females (Fisher’s exact test: P = 0.017) but not males (P = 0.524) (Fig. 4I). Among females, no weak hierarchies were observed in BALB triads, compared with 62% of C57 and 38% of *Shank3b⁻/⁻* triads, and no transitive hierarchies were observed in C57-background females. Pairwise comparisons confirmed that C57 females differed from BALB females (P = 0.003), whereas *Shank3b⁻/⁻* females did not differ significantly from either wild-type strain (BALB: P = 0.158; C57: P = 0.588). These results suggest that genetic background shapes the structure of marking hierarchies in females, with BALB females consistently forming differentiated hierarchies and C57-background females frequently forming weak ones.

#### Void Spot Summary

In summary, BALB mice of both sexes urinated more frequently in the void spot assay compared to mice on a C57 background. Most BALB males exhibited dominant urination patterns, as did nearly a quarter of BALB females. However, among mice on a C57 background, very few males showed dominant urination patterns, and almost no females did. When we compared variation in total void volume among mice in a triad, we found hierarchical structure in BALB mice of both sexes and C57 males, but weak hierarchies in C57 females and *Shank3b*⁻/⁻ mice of both sexes. This suggests that urinary scent marking reflects social hierarchy in wild-type males with intact social recognition (BALB and C57), as well as BALB females.

### Competitive performance is repeatable within assays but rarely generalizes across behavioral contexts

To determine whether competitive ability generalizes across assays, we first assessed the repeatability of individual performance within each assay, using hierarchy stability in the tube test and trial-to-trial correlations in the warm spot and void spot assays. Next, we placed performance measures from all three paradigms on a common scale. Finally, we calculated measures of cross-domain consistency and specialization to test whether high performance in one assay predicts high performance in the others.

#### Within-Assay Consistency

Across strains and sexes, triads achieved a high degree of hierarchy stability in the tube test by trial 30 (mean S index = 0.94, 95% confidence interval (CI) [0.93, 0.96]) (**Fig. 5A**). In the warm spot assay, cumulative warm spot duration was repeatable across trials (r = 0.43, 95% CI [0.31, 0.54]; Pearson’s correlation: t_187_ = 6.54, P < 0.001) (**Fig. 5B**). Similarly, total void volume was consistent across both baseline (r = 0.54, 95% CI [0.43, 0.64], t = 8.77, P < 0.001) and urine stimulus trials (r = 0.72, 95% CI [0.65, 0.78], t = 14.68, P < 0.001) (**Fig. 5C**). Together, these findings demonstrate that individual performance was repeatable in all three assays, reflecting stable differences among mice within a domain.

**Figure 5.**
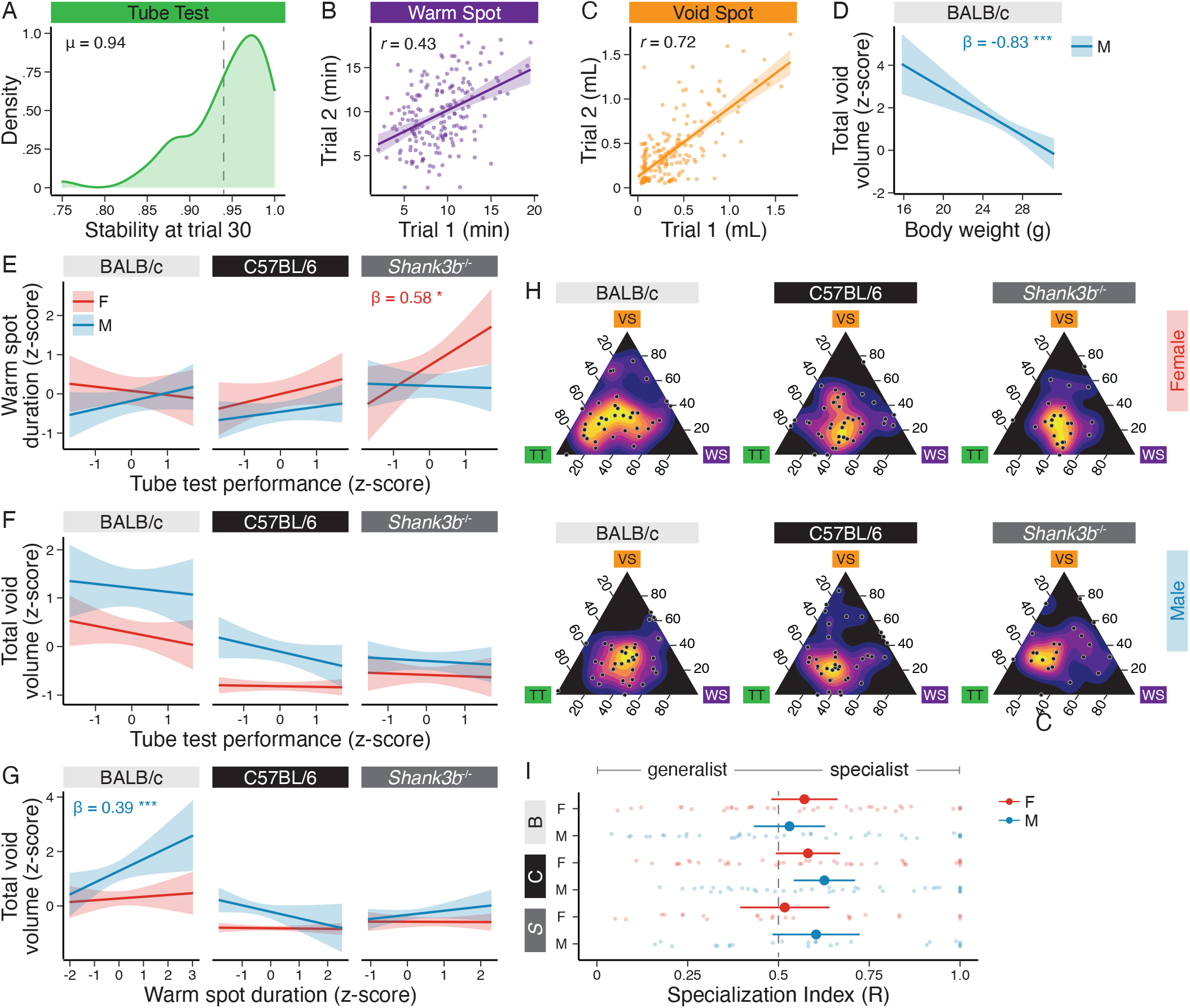
Performance comparisons within and across behavioral assays. **(A)** Distribution of hierarchy stability values (S index) at tube test trial 30. **(B)** Correlation between cumulative warm spot duration in trial 1 and trial 2. **(C)** Correlation between total void volume in trial 1 and trial 2 (urine stimulus trials). **(D)** Relationship between body weight and total void volume in BALB/c males. **(E)** Relationship between tube test performance (David’s score) and cumulative warm spot duration by strain and sex. **(F)** Relationship between tube test performance (David’s score) and total void volume by strain and sex. **(G)** Relationship between cumulative warm spot duration and total void volume by strain and sex. For (E), (F), and (G), all performance metrics were z-scored; lines indicate linear model fits and shaded areas denote 95% confidence intervals. β = standardized simple slopes for females (red) and males (blue). **(H)** Ternary plots illustrating patterns of behavioral performance across strains and sexes. The location of each point represents the relative performance across three behavioral assays for each mouse. **(I)** Behavioral specialization index (R) distributions by strain and sex.

#### Morphology

To test the prediction that larger animals have greater competitive ability, we examined the relationship between body weight and performance across all three assays. Body weight was not significantly associated with tube test performance (LMM: *t* = 1.06, *P* = 0.293) or warm spot duration (LMM: *t* = 1.15, *P* = 0.251). In BALB males only, body weight was negatively associated with total void volume (β = −0.83, 95% CI [−1.15, −0.51], *P* < 0.001), indicating that smaller males scent marked more than larger males (**Fig. 5D**).

#### Between-Assay Consistency

To determine whether dominance is a stable individual trait that generalizes across domains, we compared competitive ability across assays. Despite clear trial-to-trial stability *within* each assay, we found relatively little concordance *between* assays. Estimated marginal slopes from an LMM revealed a significant positive association between tube test performance (David’s score) and cumulative warm spot duration in *Shank3b*⁻/⁻ females only (β = 0.58, 95% CI [0.07, 1.08], P = 0.026) (**Fig. 5E**). Additionally, in BALB males, cumulative warm spot duration was positively associated with total void volume (β = 0.27, 95% CI [0.04, 0.49], P = 0.019) (**Fig. 5G**). No other slopes differed significantly from zero, suggesting that performance in one domain was generally a poor predictor of performance in another. Similarly, rank rarely generalized across assays: only 9.5% of mice occupied the same rank in all three assays (**Fig. S2**).

To compare relative performance across behavioral domains, we used ternary plots depicting the proportional contribution of each assay to an individual’s total normalized performance (Fig. 5H). In this representation (Fig. 1B), generalists, whose performance is evenly distributed across assays, cluster near the center of the plot. Conversely, individuals specializing in a single assay (single specialists) are positioned closer to a vertex, whereas those specializing in two assays (dual specialists) are positioned closer to an edge. Because the plot represents relative rather than absolute performance, an individual’s position is independent of its overall competitive success. Individuals were broadly distributed across the ternary plot, revealing substantial variation in behavioral specialization within each strain and sex.

To quantify these differences in behavioral specialization, we calculated a specialization index (R) ranging from 0 (equal relative performance across all three assays) to 1 (no contribution from at least one assay) (**Fig. S3**). Higher R values indicate increasingly uneven performance across behavioral contexts, consistent with specialization in one or two domains. We found that R varied substantially among individuals, indicating a broad continuum of behavioral specialization (mean R value = 0.57, 95% CI [0.53, 0.61]), but did not differ significantly by strain or sex (**Fig. 5I**). Altogether, these findings suggest that competitive ability is not a single, generalized trait, but rather a collection of context-dependent behaviors in which individuals exhibit varying degrees of specialization. Despite stable competitive relationships and repeatable behavior within individual assays, social rank rarely generalized across behavioral domains.

## DISCUSSION

In this study, we systematically assessed social hierarchy in laboratory mice using three established behavioral paradigms. Across these three domains, we found significant effects of sex, genetic background (BALB vs. C57), and genotype (*Shank3b*⁻/⁻ vs. C57) on both individual performance and hierarchy structure. Despite clear signatures of hierarchical organization and repeatable competitive performance within each assay, the inconsistency of rank across domains suggests that these assays capture largely independent features of competitive ability.

An important feature of our design is that we assessed hierarchies in triads instead of pairs. A triad is the smallest group in which transitivity can be assessed: each individual competes with two opponents, and the three relationships can form either a linear or a cyclic arrangement (55, 56). Triads also allowed us to quantify hierarchy structure and classify hierarchy type within each assay, and to ask whether the same individuals organize differently across behavioral contexts.

Although cyclic dominance was rare, the structure of linear hierarchies differed across assays. The tube test tended to produce transitive hierarchies, with competitive ability decreasing progressively across ranks, consistent with previous studies (16, 28, 57). In contrast, competition for warmth tended to produce despotic hierarchies, particularly among females, in which a single individual monopolized the resource, consistent with recent findings from a similar assay (58). In the void spot assay, hierarchy structure varied by strain and sex, with pronounced rank differentiation in BALB mice of both sexes and C57 males, but weaker hierarchies in C57 females and *Shank3b*⁻/⁻ mice. These results are consistent with more dominant-like urine marking behavior in BALB mice (54) and complement evidence that *Shank3* mutation can alter competitive behavior in the tube test (37). Taken together, these findings demonstrate that different behavioral contexts produce distinct patterns of hierarchical organization, even within the same group of individuals.

In the tube test, a measure of head-to-head competition, the decision to advance or retreat may reflect both self-assessment of competitive ability and memory of past wins or losses against a known opponent (59, 60). For example, mice dynamically modulate defensive responses based on the status of known competitors (16). Consistent with a role for social recognition (42–45), *Shank3b*⁻/⁻ mice of both sexes took twice as long to resolve contests as C57 controls. *Shank3b*⁻/⁻ females also showed impaired hierarchy dynamics: unlike the other strains, they lacked strong winner effects, and their hierarchies were significantly less stable over time than those of C57 controls. Moreover, no *Shank3b*⁻/⁻ female triads formed transitive or despotic hierarchies, with all classified as either weak or co-dominant. In contrast, *Shank3b⁻/⁻* males did have intact winner effects and stable hierarchies. One interpretation of these results is that males of all strains rely more heavily on self-assessment of their own competitive ability, whereas females may depend more on opponent-specific memory. The female-specific loss of winner effects and lack of a clearly differentiated dominant individual in *Shank3b*⁻/⁻ female triads suggest that social recognition may be more critical for establishing tube test hierarchies in females than in males.

In the warm spot assay, which measures group competition for access to warmth, females of all strains spent more time on the warm spot than males (e.g., 66% vs. 55% of the trial for rank 1 females and males, respectively), potentially reflecting greater thermoregulatory demand. Female mice tend to have a higher core body temperature than males (61), potentially generating a greater temperature gradient between the body and the environment that drives stronger heat-seeking behavior. Females also tended to form stronger hierarchies than males, particularly on a C57 background, with high-ranking females establishing despotic-like dominance by actively displacing other mice from the warm spot. Additionally, *Shank3b⁻/⁻* mice of both sexes spent more total time on the warm spot than either wild-type strain, and they also have higher core body temperatures than C57 controls (62), which could similarly increase heat-seeking. However, *Shank3b⁻/⁻* triads did not form stronger hierarchies than their C57 counterparts, indicating that elevated body temperature may increase motivation to occupy the warm spot without necessarily intensifying competition for it.

Finally, the void spot assay, which measures territorial marking, revealed clear strain differences in urination behavior. Specifically, BALB mice of both sexes urinated significantly more than the other two strains. The high rate of marking in BALB females was unexpected, given that urinary scent marking is generally considered a male-typical behavior (63). Nearly a quarter of BALB females displayed dominant-like urination patterns, comparable to C57 males, even though C57 is the strain most commonly used in scent marking studies (64, 65). In both sexes, mouse urine contains pheromones that signal individual identity, health, reproductive status, and social rank (66–70). Because mice advertise these signals through scent marking, it is perhaps unsurprising that BALB females would also use urinary marks to communicate social information (54).

Although C57 males marked less than BALB males, they still showed clear rank differentiation in urination behavior among triad members, whereas hierarchical organization was attenuated in *Shank3b⁻/⁻* males. Because the void spot assay is conducted individually and does not require social recognition per se, this attenuation may instead reflect altered olfactory processing or the absence of a dominance relationship established outside the assay, such as in the home cage. Void spot patterns are often interpreted as a readout of dominance established through resident-intruder encounters or co-housing (71). Notably, however, *Shank3b⁻/⁻* males formed tube test hierarchies comparable to those of C57 males, indicating that the same individuals can show intact hierarchies in one domain but attenuated hierarchies in another. Together, these findings reveal substantial variation in urination patterns and their hierarchical organization across strains and sexes and further illustrate that hierarchical organization in one behavioral context need not predict organization in another.

Across all three behavioral paradigms, individual performance was repeatable within each assay, indicating that mice display stable competitive behavior within a specific behavioral context. Despite this within-assay consistency, individual competitive performance did not readily generalize across behavioral domains. With two exceptions (positive correlation between tube test and warm spot performance in *Shank3b⁻/⁻* females and between warm spot and void spot performance in BALB males), competitive ability was largely independent across assays. These findings suggest that each assay captures distinct features of competitive ability, with individuals exhibiting varying degrees of specialization across behavioral domains.

These results add to growing evidence that competitive success in laboratory assays is domain-specific. Social dominance is traditionally defined by priority of access to food, mates, and territory, yet competitive success need not be consistent across domains (4, 29). Indeed, in male mice, tube test rankings can diverge from rankings based on resource competition, agonistic interactions, and territorial marking (39, 40). Here, we extend these observations to females, another genetic background (BALB), and a genetic model of impaired social recognition (*Shank3b*⁻/⁻), and additionally show that the same triads organize into different hierarchy types across domains. Moreover, because performance was repeatable within each assay, this independence is unlikely to reflect measurement noise. This does not diminish the utility of these assays, but it does change how they should be interpreted: rather than interchangeable measures of a single, generalized dominance trait, they are better understood as task-specific assays that capture distinct features of competitive behavior (72).

Altogether, our study demonstrates the value of assessing social hierarchy across multiple behavioral contexts, revealing that competitive ability is multidimensional and that different assays capture complementary aspects of social behavior. These findings have implications for neuroscience studies seeking to understand the mechanistic basis of social behavior in the laboratory mouse, as neural mechanisms identified using a single assay may support domain-specific rather than generalized competitive success.

---

**Figure S1.**
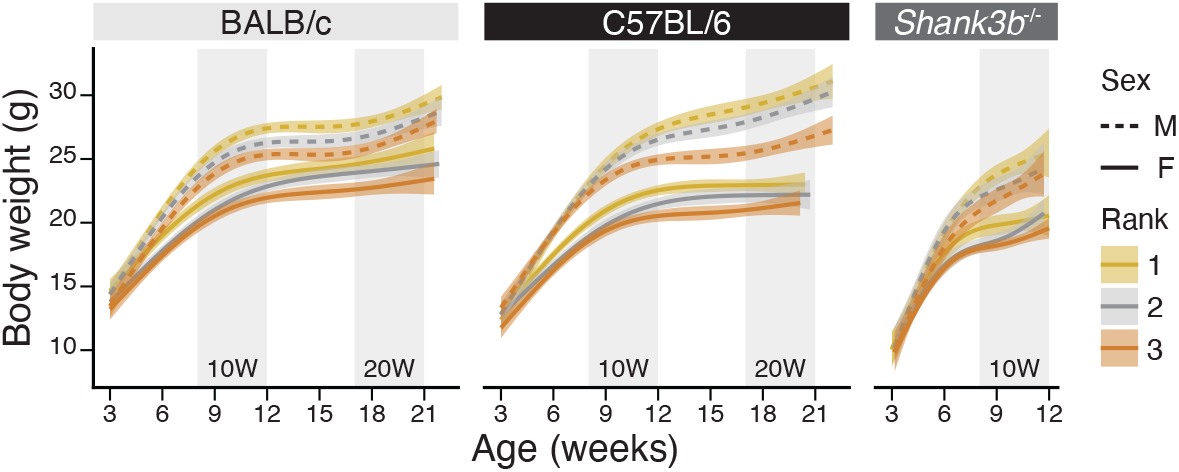
Body weight trajectories by strain and sex. Lines indicate generalized additive model (GAM)-smoothed trajectories fitted by rank (rank 1 = heaviest, rank 3 = lightest) for BALB/c, C57BL/6, and *Shank3b*^-/-^ mice. Dashed lines represent males and solid lines represent females. Shaded areas denote 95% confidence intervals. Grey vertical bars show the testing period for 10-week-old (weeks 8–12) and 20-week-old (weeks 17–21) triads.

**Figure S2.**
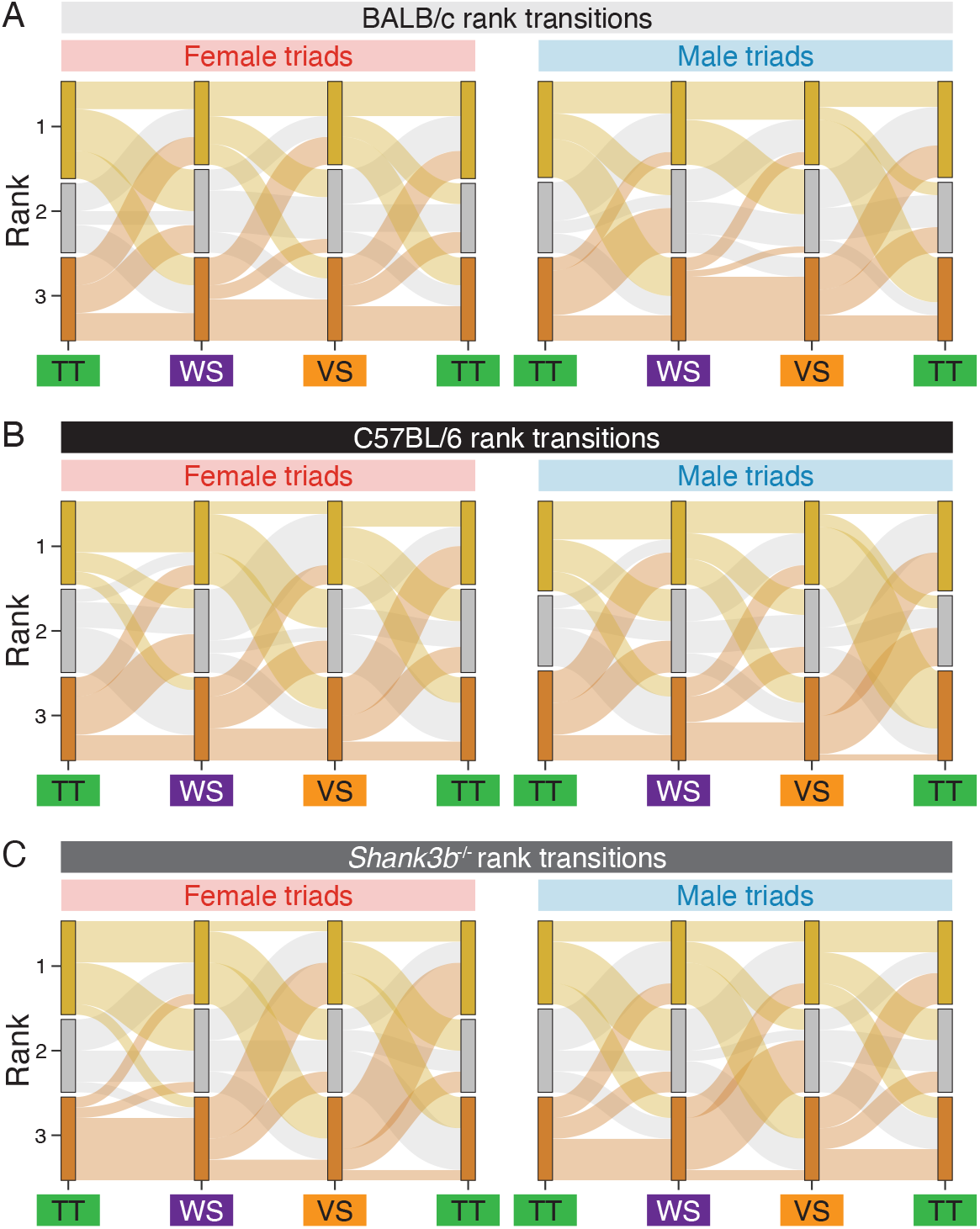
Rank concordance across behavioral assays. Each Sankey diagram tracks individual rank assignments from the tube test (TT) to the warm spot assay (WS) to the void spot assay (VS), shown separately for **(A)** BALB/c, **(B)** C57BL/6, and **(C)** *Shank3b*^-/-^ mice. Within each assay, mice were ranked 1–3 based on David’s score, mean cumulative warm spot duration, or mean total void volume, where rank 1 indicates the highest-ranking individual. Flow width is proportional to the number of mice following each rank trajectory.

**Figure S3.**
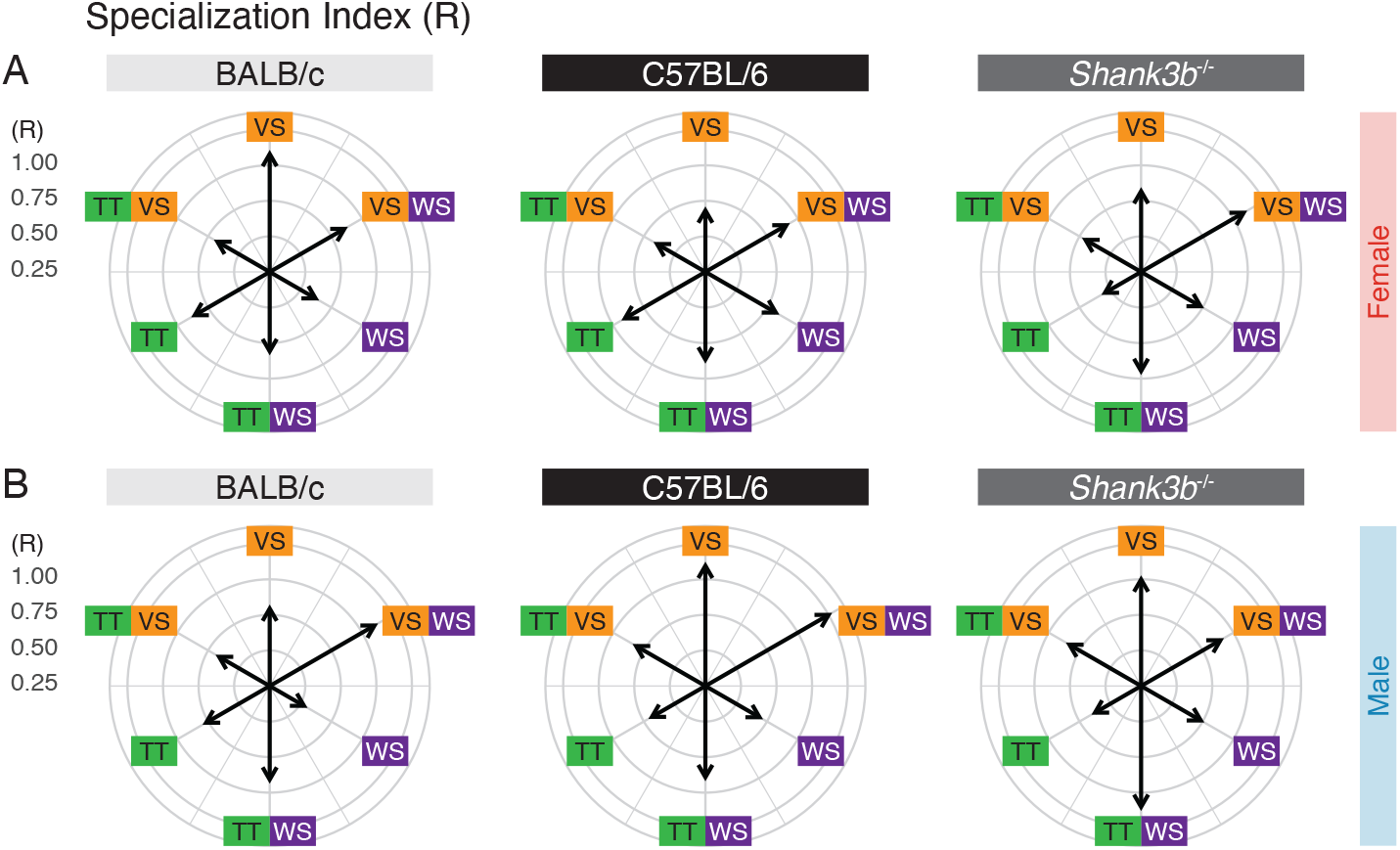
Competitive specialization across behavioral assays. Radial plots show the median specialization index (R) for mice exhibiting specialization in one (TT, WS, VS) or two (TT/WS, TT/VS, WS/VS) behavioral assays, shown separately for **(A)** females and **(B)** males. Arrow direction indicates the behavioral domain(s) of specialization, and arrow length represents the median R value, ranging from 0 (equal relative performance across all three assays) to 1 (no contribution from at least one assay). Higher R values indicate increasingly uneven performance across behavioral contexts. Abbreviations: TT = tube test; WS = warm spot; VS = void spot.

