## Extended Methods for "Social hierarchy assays measure independent features of competitive ability"

#### Animals

BALB/c (stock no. 000651), C57BL/6 (stock no. 000664), and Shank3B$^{-}$ (stock no. 017688) mice were obtained from The Jackson Laboratory (Bar Harbor, ME) and maintained in the Bedford Lab at the University of Wyoming (UW). *Shank3b*$^{+/-}$ mice were backcrossed onto a C57BL/6 background for a minimum of 6 generations and subsequently maintained as homozygous knockouts (*Shank3b*$^{-/-}$). A parallel *Shank3b*$^{+/+}$ control line was generated from *Shank3b*$^{+/-}$ intercrosses and maintained alongside the *Shank3b*$^{-/-}$ line. Because no behavioral differences were detected between *Shank3b*$^{+/+}$ mice and wild-type C57BL/6 mice, these groups were combined for analysis. A total of 204 animals (68 triads) were used in this study (**Table S1**). All experimental protocols were approved by the UW Institutional Animal Care and Use Committee (protocol no. 2022-0126).

**Table S1.** Triad sample sizes.

| Background | Strain | Sex | Test Age | Source | No. Triads | Total |
|---|---|---|---|---|---|---|
| BALB/c | BALB/c | F | 10 weeks | JAX | 2 | 12 |
| | BALB/c | F | 10 weeks | UW | 4 | |
| | BALB/c | F | 20 weeks | UW | 6 | |
| | BALB/c | M | 10 weeks | JAX | 2 | 14 |
| | BALB/c | M | 10 weeks | UW | 5 | |
| | BALB/c | M | 20 weeks | UW | 7 | |
| C57BL/6 | *Shank3b*$^{+/+}$ | F | 10 weeks | UW | 3 | 13 |
| | C57BL/6 | F | 10 weeks | JAX | 2 | |
| | C57BL/6 | F | 10 weeks | UW | 3 | |
| | C57BL/6 | F | 20 weeks | UW | 5 | |
| | *Shank3b*$^{+/+}$ | M | 10 weeks | UW | 3 | 13 |
| | C57BL/6 | M | 10 weeks | JAX | 2 | |
| | C57BL/6 | M | 10 weeks | UW | 3 | |
| | C57BL/6 | M | 20 weeks | UW | 5 | |
| | *Shank3b*$^{-/-}$ | F | 10 weeks | UW | 8 | 8 |
| | *Shank3b*$^{-/-}$ | M | 10 weeks | UW | 8 | 8 |
| | | | | | | **68** |

**Husbandry.** Animals were housed under a 12:12 h reversed light–dark cycle at 22°C in standard polycarbonate cages (7½” W × 11½” L × 5” H; Ancare). Bedding consisted of Rocky Mountain softwood screened shavings and quarter-inch corncob bedding (Inotiv). Enrichment included a cotton nestlet (Ancare) and a red polycarbonate hut (Bio-Serv). Water and Mouse Diet 9F (LabDiet) were provided *ad libitum*. All behavioral assays were conducted during the dark phase.

**Triads.** Age- and weight-matched mice from different litters were weaned simultaneously into same-sex groups of three (triads) at postnatal day 24 (P24). When necessary, weaning was advanced or delayed by 1–2 days so that all members of a triad could be placed into the same cage on the same day. Within triads, the mean age difference at weaning was 1.6 d and the mean weight difference was 1.3 g. We found no significant effect of wean weight on tube test performance (t = 1.00, P = 0.321), warm spot duration (t = 0.68, P = 0.499), or void volume (t = 1.06, P = 0.291). Additional C57BL/6 and BALB/c mice were obtained from The Jackson Laboratory (JAX) and co-housed in same-sex triads at 3 weeks of age. These mice were likely littermates, but because no significant behavioral differences were detected between JAX triads and those born at UW, these groups were combined for analysis (**Table S1**).

Mice were weighed at weaning and every 7 d thereafter for the duration of the experiment (**Fig. S1**). Triads were maintained undisturbed, except for weekly weighing and biweekly cage changes, for a minimum of 5 weeks. Tube test training began at 8–9 weeks of age for 10-week triads and at 17–18 weeks of age for 20-week triads. Following a 3-week testing period—encompassing the tube test, warm spot assay, and void spot assay—animals were euthanized and tissues were collected at 11–12 weeks of age (10-week triads) or 20–21 weeks of age (20-week triads). Age groups were pooled for analysis, with test age (10-week or 20-week) included as a covariate in all statistical models.

**Urine Collection.** We collected urine from adult, virgin C57BL/6 mice of both sexes from the Bedford Lab colony. Fresh urine from several mice of the same sex was pooled by pipetting and frozen in 60 µL aliquots at −20°C. Individual aliquots were thawed and warmed to 37°C in a water bath prior to use as an olfactory stimulus in the void spot assay (see below).

### Behavioral Tests

**Tube Test.** For tube test training and testing, mice were singly housed for 30 min prior to handling. Each mouse underwent 4 d of training to learn to fully traverse the tube before formal testing. Training runs were conducted in alternating directions: right-to-left (RL) and left-to-right (LR). On training days 1–2, mice completed RL (6 runs) and LR (6 runs), followed by a rest period and an additional RL (3 runs) and LR (3 runs). On training days 3–4, mice completed RL (3 runs) and LR (3 runs), followed by an additional RL (1 run) and LR (1 run), for a total of 26 training runs in each direction. If a mouse stopped in the tube, it was gently encouraged to continue using a padded wooden dowel.

All mice were trained and tested using an 18-inch (45.7 cm) clear extruded acrylic tube (United States Plastic Corp.). C57BL/6 and *Shank3b*$^{-}$/$^{-}$ mice were tested in a tube with a 1-inch (2.54 cm) inner diameter (ID), whereas BALB/c mice, which are slightly larger, were tested in a tube with a 1⅛-inch (2.86 cm) ID. Following training, each triad underwent a 30-trial round-robin tournament in which each mouse encountered the other two triad members 10 times over 5 d (20 trials per individual). Starting sides were alternated for each pair to ensure that mice encountered their opponent from both ends of the tube.

**Warm Spot Assay.** All three mice were simultaneously placed on a 19.5 cm diameter cold plate maintained at 15°C (Ugo Basile). A custom-built 4.5 cm × 5 cm platform (the “warm spot”) adjacent to the cold plate was maintained at 25°C. Assays lasted 20 min and were video recorded under infrared illumination. Individual mice were identified by unique tail markings, and each triad was tested twice. Warm spot bouts were quantified by researchers blinded to experimental condition using The Observer XT software (Noldus) to extract the number of bouts, median bout duration, and total time spent on the warm spot. To assess inter-observer consistency, a random subset of 14 trials was independently scored by two observers. Consistency scores ranged from 0.62 to 0.76, and values from independently scored trials were averaged across observers prior to analysis.

**Void Spot Assay.** Individual mice were placed in a 20.3 cm × 25.4 cm open-field arena lined with cellulose chromatography paper (Fisherbrand) and allowed to freely explore for 20 min. Two baseline trials were conducted as described above, followed by two trials in which 60 µL of same-sex urine was pipetted onto the center of the filter paper immediately prior to testing. Filter papers were imaged using a FLIR USB 3.0 camera (FL3-U3-13Y3M-C) with blue light excitation (455 nm, 1150 mW mounted LED; ThorLabs) and a GFP filter set with excitation at 460 ± 30 nm (ET460/30x) and emission at 525 ± 50 nm (ET525/50m; Chroma). Images were acquired using SpinView software (Teledyne) and quantified with custom ImageJ scripts to extract the number of void spots, mean and maximum spot volume, total void volume, and the proportion of spots and total volume in the center versus the edge of the filter paper.

### Statistics

All statistical analyses were performed in R version 4.6.0 (2026-04-24). Raw data and R scripts are available at https://github.com/bedford-lab/hierarchy.

**Tube Test.** Trial duration was analyzed using a linear mixed-effects model (LMM; function lmer, package lme4) with strain, sex, age, and the strain × sex interaction as fixed effects and pair ID as a random intercept. Post hoc pairwise comparisons were conducted using estimated marginal means (function emmeans, package emmeans) with Tukey adjustment for multiple comparisons. For each opponent pair, we calculated the proportion of trials resolved by the winner advancing rather than the loser retreating and analyzed this using a linear model (LM; function lm, package stats) with strain, sex, age, and the strain × sex interaction as predictors. Pairwise differences between strains and sexes were evaluated using emmeans.

Winner effects were modeled using a Bayesian generalized linear mixed-effects model (function brm, package brms). Outcome (win/loss) was modeled with a Bernoulli distribution and logit link as a function of current win streak, strain, sex, and their two- and three-way interactions, with random intercepts for mouse ID and partner ID. Weakly informative priors were specified as follows: Normal(0, 1.5) for the intercept, Normal(0, 1.0) for fixed-effect coefficients, and Exponential(1) for random-effect standard deviations. Posterior effects were reported as medians with 95% credible intervals (CrIs). Tukey-adjusted post hoc comparisons of strain- and sex-specific associations were conducted using estimated marginal slopes (function emtrends, package emmeans). Maximum win streak length distributions were compared across strains and between sexes using Anderson–Darling tests (function ad.test, package kSamples).

Elo ratings were calculated from the ordered sequence of contests within each triad (function elo.seq, package EloRating). Hierarchy stability was quantified using the S index (function stab_elo, package EloRating) and modeled using an LMM with trial number, strain, sex, and their two- and three-way interactions as fixed effects. Triad age (10-week or 20-week) was included as a fixed covariate, and triad ID was included as a random intercept to account for repeated measures across trials. Tukey-adjusted post hoc comparisons were conducted using estimated marginal slopes (emtrends). Additionally, S index values were extracted at trials 10, 20, and 30 and compared across strains and between sexes using LMs with strain, sex, age, and the strain × sex interaction as predictors; pairwise differences were evaluated using emmeans.

David's scores (DS) were calculated from the win/loss interaction matrix for each triad (function DS, package EloRating), and pairwise dominance separations were computed as follows: the rank 1 to rank 2 difference ($\Delta_{12} = DS_1 - DS_2$), the rank 2 to rank 3 difference ($\Delta_{23} = DS_2 - DS_3$), and the overall rank 1 to rank 3 difference ($\Delta_{13} = DS_1 - DS_3$). Hierarchy type was classified using the empirical distribution of $\Delta_{13}$ values across all triads: those with $\Delta_{13}$ below the 25th percentile were classified as weak (egalitarian), and the remainder were classified as hierarchical. Hierarchical triads were subcategorized based on the position of the rank 2 individual relative to ranks 1 and 3, dividing the rank 1 to rank 3 interval into three equal segments. If $DS_2$ fell within the upper third (closer to $DS_1$), the hierarchy was classified as co-dominant; if $DS_2$ fell within the middle third, it was classified as transitive; and if $DS_2$ fell within the lower third (closer to $DS_3$), it was classified as despotic. The distribution of hierarchy types across strains was compared separately in males and females using Fisher's exact tests (function fisher.test, package stats).

**Warm Spot Assay.** Cumulative warm spot duration was analyzed using an LMM with rank, strain, sex, and their two- and three-way interactions as fixed effects. Age was included as a fixed covariate, and mouse ID was included as a random intercept to account for repeated measures across trials. Marginal effects of strain, sex, and their interaction were estimated using emmeans with Tukey adjustment for multiple comparisons. The number of bouts per trial was analyzed using an LMM with strain, sex, age, and the strain × sex interaction as fixed effects and mouse ID as a random intercept. Median bout duration across both trials was analyzed using an LM with strain, sex, age, and the strain × sex interaction as predictors. Total displacements per trial was analyzed using an LMM with strain, sex, age, and the strain × sex interaction as fixed effects and triad ID as a random intercept to account for non-independence among mice in the same triad. For bout number, bout duration, and displacements, marginal effects of strain and the strain × sex interaction were estimated using emmeans with Tukey adjustment for multiple comparisons.

Directed network graphs were constructed for each trial with individual mice as vertices and displacement counts as weighted directed edges (function graph_from_data_frame, package igraph). We extracted vertex-level statistics including out-strength (number of displacements initiated), in-strength (number of displacements received), and net strength (out-strength minus in-strength; function strength, package igraph). Network-level statistics included out-centralization, in-centralization, and a linearity index reflecting the proportion of interactions occurring in the direction of the triad rank order.

Individuals within each triad were ranked by mean cumulative warm spot duration, and net displacement values were compared across ranks using an LM with rank, strain, sex, and their two- and three-way interactions as predictors, with age included as a covariate. Pairwise rank differences within each strain and sex were evaluated using emmeans.

Hierarchy type was classified based on pairwise differences in mean warm spot duration among rank 1, rank 2, and rank 3 individuals within each triad, following the same criteria described above for the tube test. The distribution of hierarchy types between males and females was compared using Fisher's exact test.

**Void Spot Assay.** The probability of voiding was modeled using a generalized linear mixed-effects model (GLMM; function glmer, package lme4) with a binomial error distribution and logit link, including strain, sex, age, and stimulus as fixed effects and mouse ID as a random intercept. Pairwise differences between strains and stimulus conditions were assessed using emmeans. Total void volume per trial was $\log_{10}$-transformed and analyzed using an LMM with strain, sex, stimulus, and all two-way interactions as fixed effects, with age as a covariate and mouse ID as a random intercept. Marginal effects of strain and the strain × sex interaction were estimated using emmeans with Tukey adjustment for multiple comparisons.

Urination patterns were classified using linear discriminant analysis (LDA; function lda, package MASS). Three researchers blinded to experimental condition independently classified 175 randomly selected filter papers as either dominant, subordinate, or neutral. Dominant patterns consisted of many small-volume voids distributed throughout the arena; subordinate patterns consisted of one or two large-volume voids localized to the corners; and neutral patterns consisted of a small number of intermediate-volume voids distributed throughout the arena. The three observers reached unanimous agreement on 142 filter papers, which were used as the training set. LDA models were trained on six quantitative features: number of void spots, mean and maximum spot volume, total void volume, and the proportion of spots and total volume in the center versus the edge of the arena. Only trials with at least one void were included, and all predictor variables were z-scored prior to analysis. A robust LDA model was used for final classification. Model performance was assessed by resubstitution accuracy and leave-one-out cross-validation. The model achieved 93.0% resubstitution accuracy (95% CI [0.87, 0.97], $\kappa = 0.881$) and 89.4% leave-one-out cross-validated accuracy (95% CI [0.83, 0.94], $\kappa = 0.824$). Cross-validated sensitivities were 91%, 87%, and 89% for dominant, subordinate, and neutral classes, respectively.

A scent marking index (SMI) was calculated as: $SMI = \max(LD1) - LD1 + 1$, such that higher values indicate more dominant-like urination patterns. SMI values were analyzed using an LMM with strain, sex, stimulus, age, and the strain × sex interaction as fixed effects and mouse ID as a random intercept. Individuals were also ranked within each triad by total void volume (averaged across two urine stimulus trials). Total void volume was then compared across ranks using an LM with rank, strain, sex, and all two- and three-way interactions as predictors, with age included as a covariate. Pairwise rank differences within each strain and sex were evaluated using emmeans. The distribution of urination pattern classes (dominant, subordinate, or neutral) and hierarchy types across strains and sexes were compared using Fisher's exact tests.

**Assay Comparisons.** Trial-to-trial consistency within the warm spot and void spot assays was evaluated using Pearson's product-moment correlations (function cor.test, package stats). To compare performance across assays, three summary metrics were extracted for each mouse: David's score from the tube test, cumulative warm spot duration (mean of trials 1 and 2), and total void volume (mean of trials 1 and 2). All three metrics were z-scored prior to analysis. Pairwise associations between performance metrics were examined using LMMs with strain, sex, and their interaction as fixed effects, triad age as a fixed covariate, and triad ID as a random intercept. Tukey-adjusted post hoc comparisons of strain- and sex-specific associations were conducted using estimated marginal slopes (emtrends).

To assess relative performance across behavioral domains, each metric was min–max normalized to a 0–1 scale within strain and sex. For each mouse, normalized values were summed to calculate a total dominance score, and each assay's contribution was expressed as a proportion of this total. The three resulting proportions summed to one and were visualized using ternary plots (function ggtern, package ggtern), with kernel density estimates overlaid to illustrate the distribution of individuals within ternary space.

To visualize rank consistency across assays, mice were ranked within each triad based on David's score from the tube test, mean cumulative warm spot duration, and mean total void volume. Rank transitions between assays were visualized using Sankey diagrams (function sankeyNetwork, package networkD3) for each strain and sex. Diagrams displayed rank transitions from assay to assay, with link widths proportional to the number of mice exhibiting each transition (**Fig. S2**).

Behavioral specialization was quantified using an index (R) calculated as $R = 1 - 3 \times \min(TT, WS, VS)$, where TT, WS, and VS represent the proportional contributions of each assay to total normalized performance. R ranges from 0 (indicating equal contributions across all three assays) to 1 (indicating no contribution from at least one assay). Higher R values indicate greater specialization in one or two behavioral contexts. To characterize the direction of specialization, individuals were assigned to one of six reference strategies representing single-assay (TT, WS, or VS) or dual-assay (TT/WS, TT/VS, or WS/VS) specialization. Strategies were assigned based on the minimum Euclidean distance between each individual's position in ternary space and six reference trajectories extending from the center toward each vertex or edge midpoint. Median R was calculated for each strategy within each strain and sex and visualized using radial plots, with vector direction indicating specialization strategy and vector length indicating median R (**Fig. S3**). Differences in R across experimental groups were analyzed using an LM with strain, sex, and the strain × sex interaction as predictors. Type III tests (function Anova, package car) were used to evaluate the significance of each effect.

**Morphology.** Associations between body weight and z-scored performance metrics were tested using LMMs with strain, sex, age, and the strain × sex interaction as fixed effects and triad ID as a random intercept.
